# Crystal structures of two tandem malectin-like receptor kinases involved in plant reproduction

**DOI:** 10.1101/251959

**Authors:** Steven Moussu, Sebastian Augustin, Andra Octavia Roman, Caroline Broyart, Julia Santiago

## Abstract

A complex cell-to-cell communication between the male pollen tube and the female reproductive organs is required for plant fertilization. A family of *Catharanthus roseus* Receptor Kinase-1 ( *Cr*RLK1L) membrane receptors has been genetically implicated in this process. Here we present crystal structures of the CrRLK1Ls ANXUR1 and 2 at 1.48 and 1.1 Å resolution, respectively. Our structures reveal a novel arrangement of two malectin-like domains connected by a short βhairpin linker and stabilized by calcium ions. The canonical carbohydrate interaction surfaces of related animal and bacterial carbohydrate binding modules are not conserved among plant CrRLK1Ls. In line with this, we failed to detect binding of chemically diverse oligosaccharides to ANXUR1 and HERCULES1. Instead, CrRLK1Ls have evolved a protein-protein interface between their malectin domains, which forms a deep cleft lined by highly conserved aromatic and polar residues. Analysis of the glycosylation pattern of different CrRLK1Ls and their oligomeric states together suggests that this cleft could resemble a binding site for a ligand required for *Cr*RLK1Ls receptor activation.

## INTRODUCTION

Plant cells are surrounded by a dynamic, carbohydrate-rich cell wall, which is constantly remodeled to enable coordinated growth and development, and serves as a link with the outside world (1). To sense and integrate environmental and internal signals, plants have evolved a set of membrane receptor kinases (RKs), whose extracellular domains are facing the wall compartment (2, 3). Key players in monitoring the cell wall status and regulating cell expansion are members of the plant-specific CrRLK1 family (4–6). From the 17 members in Arabidopsis, 10 have been functionally characterized: FERONIA (FER), ANXUR1 (ANX1), ANXUR2 (ANX2), BUDDHA PAPER SEAL 1/2 (BUPS1/2), HERCULES1 (HERK1), HERCULES2 (HERK2), Ca^2+^cyt-associated protein kinase 1 (CAP1/ERULUS), THESEUS1 (THE1) and CURVY1 (CVY1) (7). FER, ANX1/2 and BUPS1/2 have been involved in cell communication events regulating plant fertilization, with FER acting in the female gametophyte and ANX1/2 and BUPS1/2 in the male gametophyte, the pollen. Loss of FER function impairs pollen tube reception (8), whereas ANX1/2 together with BUPS1/2 help maintaining pollen tube integrity during polarized tip growth, assuring fertilization (9–11). FER, THE1, HERK1 and HERK2 have been implicated in regulating cell expansion during vegetative growth (12, 13), with FER having additional roles in plant immunity (14–16). THE1 has been described to be able to sense structural changes of the cell wall and to regulate lignin accumulation in cellulose-deficient mutants (17). Recent studies have also linked a member of this receptor family with the control of cell morphogenesis and cytoskeleton assembly (18).

CrRKL1 family members localize to the plasma membrane (17) and are composed of a cytoplasmic kinase domain, a single transmembrane helix and a variable ligand-binding ectodomain. The extracellular domain shows weak sequence homology to animal malectin carbohydrate binding domains (6, 19). A domain swap analysis of several members of the family suggested that the signaling specificity of *Cr*RLK1Ls is encoded in their extracellular domain. In contrast, their intracellular kinase domain can be interchanged, suggesting that this receptor family may be sharing common downstream signaling components (20). In the case of FER, the connecting transmembrane domain has also been involved in regulating the receptor activation (21). Whether the extracellular domains of CrRLK1Ls actually form ligand binding domains for carbohydrate and/or protein ligands still remains to be characterized at the molecular level. Thus far, several secreted ∼40 amino-acid peptides of the RAPID ALKALINIZATION FACTORs (RALFs) family have been proposed as ligands for FER, ANX1/2 and BUPS1/2 (10, 16, 22, 22), but CrRLK1Ls have also been speculated to interact directly with cell wall components (4, 7, 23). Biochemical analysis of CrRLK1Ls has been hampered by the difficulties to produce active, recombinant protein samples on one hand, and by the overwhelming chemical complexity of the plant cell wall on the other hand. Here we report the expression, purification and biochemical and crystallographic characterization of different CrRLK1Ls as a first step to understand cell wall sensing and signaling mechanisms in plants.

## Results

We produced the ectodomains of Arabidopsis ANX1, ANX2, HERK1 and THE1 by secreted expression in insect cells and purified the N-glycosylated proteins to homogeneity (see Methods). We obtained crystals for all receptors and diffraction quality crystals for ANX1 and 2. We determined the structure of ANX1 via single isomorphous replacement using a platinum derivative and the structure of ANX2 by molecular replacement using the ANX1 structure as search model (Table 1). The structures of ANX1 and ANX2 were refined to 1.48 Å and 1.1 Å resolution, respectively, with residues 26-411 being well-defined by electron density. The ectodomains of ANX1 and ANX2 superimpose with an r.m.s.d of ∼0.6 Å, comparing 375 corresponding C_α_ atoms in their ectodomains. ANX1 folds into two individual βsandwich malectin-like domains consisting of 4 antiparallel β-strands, connected by long loops and short helical segments (Fig. 1*A*). The two domains tightly pack against each other, placing their βsandwich cores at an angle of ∼ 85° (Fig. 1*A*). A β-hairpin linker connects the N-terminal (mal-N) and the C-terminal (mal-C) malectin domains. The two malectin-like domains share an extensive hydrophilic interface with each other and with the β-hairpin linker (∼850 Å^2^ buried surface area) (Fig. 1*A*). Mal-N and mal-C domains align closely (root mean square deviation r.m.s.d. is ∼ 1.5 Å comparing 138 corresponding C_α_ atoms) (Fig. 1*B*). The two highly conserved cysteine residues in CrRLK1Ls are part of the structural cores of the mal-N and mal-C β-sandwich domains rather than being engaged in a disulfide bond (S-S distance is ∼38.4 Å) (Fig. 5*C*). A search with the program DALI (24) returned the animal malectin from *Xenopus laevis* as the closest structural homolog of ANX1 (DALI Z-score is 13, r.m.s.d. is ∼2.2 Å comparing 127 corresponding C_α_ atoms; Fig. 1*C*) (19). In addition, the ANX1 malectin domains share structural features with other carbohydrate binding modules (CBMs), such as CBM22 and CBM35 from bacterial xylanases and hydrolases involved in plant cell wall degradation (Figs. 1*D*, 2) (25, 26). In mal-N and mal-C from ANX1 and ANX2, we located two calcium binding sites, which appears to function in structural stabilization rather than acting as an enzymatic cofactor (Fig. 2*A, B*). Similar calcium binding sites have been previously described for other CBMs (Fig. 2*C*) (27).

**Figure 1.**
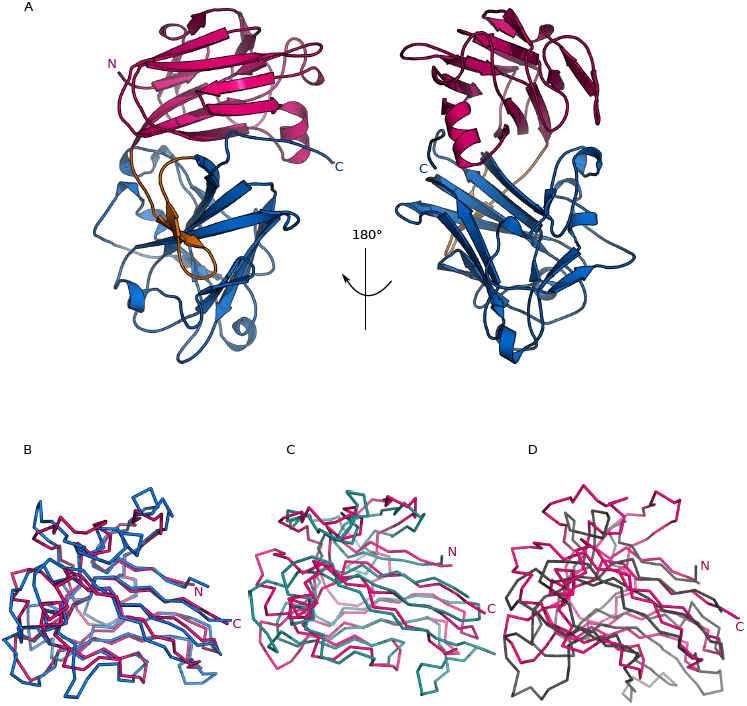
Architecture of the tandem malectin-like ectodomain of ANXUR1. *A*, front and 180° rotated views of the ANX1 ectdomain (ribbon diagram) with mal-N shown in magenta, the β-hairpin in orange and mal-C in blue. *B*, structural superposition of the ANX1 mal-N domain (C_α_-trace, in magenta) onto mal-C(blue). Root mean square deviation r.m.s.d. is ∼2 Å comparing 127 corresponding C_α_ atoms. *C* and *D*, structural superpositions of the ANX1 mal-N (magenta) onto the (*C*) animal malectin from *Xenopus laevis* (light-blue) (PDB-ID: 2K46) (19) and (*D*) the bacterial carbohydrate binding module (CBM22) from *Paenibacillus barcinonensis* (shown in gray) (PDB-ID: 4XUR) (25). R.m.s.d.’s are ∼2.2 Å comparing 127 corresponding C_α_ atoms, and ∼3.1 Å comparing 119 corresponding C_α_ atoms, respectively.

**Figure 2.**
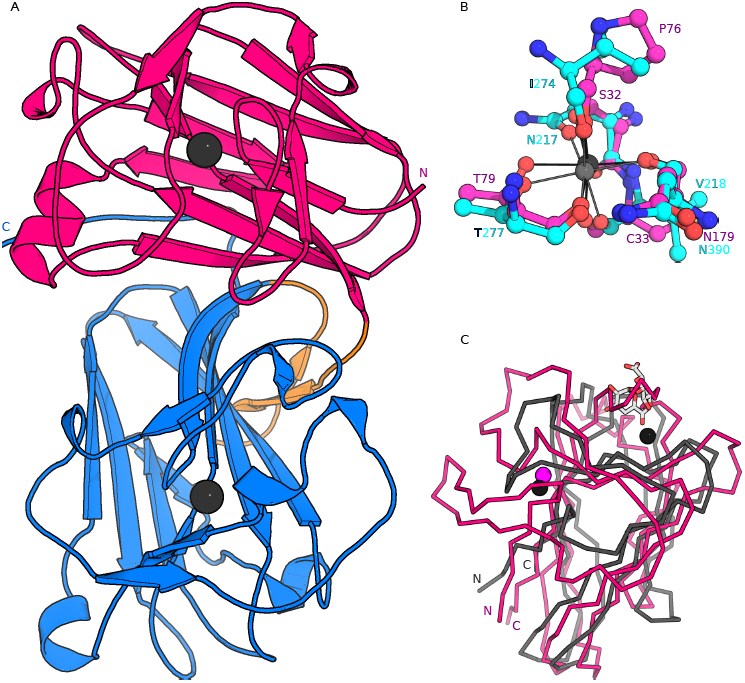
The ANX1 malectin-like domains are structurally stabilized by calcium ions. *A*, ribbon diagram of the ANX1 ectodomain colored as in Fig. 1 and including the positions of two stabilizing calcium-ions (black spheres). *B*, details of the ANX1 mal-N (magenta) and mal-C (cyan) calcium ion-binding sites. Residues are in bonds representation, metal coordinating interactions are indicated by solid lines. *C*, the ANX1 calcium ion positions map to structural not enzymatic calcium binding sites in CBMs. Structural superposition of the ANX1 mal-N domain (ribbon diagram, in magenta) with the CBM35 bound to digalacturonic acid (PDB: 2VZQ; r.m.s.d is ∼3.7 Å comparing 68 corresponding C_α_ atoms) (26) shown in gray. CBM35 contains two calcium ions, depicted as black spheres, one in its catalytic binding site in close proximity to the carbohydrate, and another one having a structural role located at the opposite face of the binding groove. The position of the latter one corresponds to the observed calcium binding site in ANX1 (shown as a magenta sphere).

**Table 1.** Crystallographic data collection and refinement statistics.

|  | **Anxur1**<br>(NH<sub>4</sub>)<sub>2</sub>PtCl<sub>4</sub><br>derivative | **Anxur1**<br>native1 | **Anxur1**<br>native 2 | **Anxur2**<br>native |
| --- | --- | --- | --- | --- |
| **Data collection** |  |  |  |  |
| Space group | *P*1 | *P*1 | *P*1 | *C*2 |
| Cell dimensions<sup>‡</sup><br>*a*, *b*, *c* (Å) | 54.78, 68.55, 69.89 | 54.23, 68.50, 69.89 | 53.98, 68.74, 70.02 | 123.56,<br>41.76,<br>93.37 |
| $\alpha$, $\beta$, $\gamma$ (°) | 88.04, 75.57, 72.29 | 88.86, 74.71, 71.93 | 88.56, 74.97, 71.74 | 90, 117.4, 90 |
| Resolution (Å) | 48.39 – 2.13<br>(2.18 – 2.13) | 48.39 – 1.89 (2.00-1.89) | 47.27 – 1.48 (1.52-1.48) | 44.29 – 1.08 (1.12 –<br>1.08) |
| *R*<sub>meas</sub><sup>#</sup> | 0.082 (1.41) | 0.112 (1.16) | 0.062 (2.17) | 0.041 (1.54) |
| CC(1/2)<sup>#</sup> (%) | 100 (67.6) | 100 (50.5) | 100 (68.0) | 100(47.1) |
| *I*/$\sigma$ *I*<sup>#</sup> | 10.7 (1.0) | 6.9 (1.0) | 17.9 (1.2) | 18.98(0.92) |
| Completeness (%)<sup>#</sup> | 96.2 (81.5) | 93.8 (93.4) | 98.4 (95.6) | 96.1 (80.54) |
| Redundancy<sup>#</sup> | 3.5 (3.3) | 1.8 (1.7) | 10.3 (10.2) | 6.5 (5.0) |
| Wilson B-factor<sup>#</sup> | 51.0 | 35.8 | 31.4 | 14.70 |
| **Phasing** |  |  |  |  |
| Resolution (Å)<sup>†</sup> | 48.39 – 3.0 |  |  |  |
| No. of sites<sup>†</sup> | 10 |  |  |  |
| Phasing power<br>(iso)<sup>†</sup> | 0.449 |  |  |  |
| Phasing power<br>(ano)<sup>†</sup> | 0.854 |  |  |  |
| FOM<sup>†</sup> | 0.344 |  |  |  |
| **Refinement** |  |  |  |  |
| Resolution (Å) |  |  | 47.27 – 1.48<br>(1.52 – 1.48) | 44.29 – 1.08 (1.11 –<br>1.08) |
| No. reflections |  |  | 14,3968 (10,394) | 165,089 (9,560) |
| *R*<sub>work</sub>/*R*<sub>free</sub><sup>\$</sup> | | | 0.172/0.192<br>(0.303/0.309) | 0.159/0.182<br>(0.470/0.535) |
| No. atoms |  |  |  |  |
| Protein |  |  | 6,130 | 3,028 |
| Glycan |  |  | 164 | 95 |
| Calcium |  |  | 4 | 2 |
| Water |  |  | 743 | 433 |
| Res. B-factors<sup>\$</sup> | | | | |
| Protein |  |  | 32.6 | 22.2 |
| Glycan |  |  | 51.2 | 37.7 |
| Calcium |  |  | 34.8 | 14.2 |
| Water |  |  | 41.8 | 39.3 |
| R.m.s deviations<sup>\$</sup> | | | | |
| Bond lengths (Å) |  |  | 0.01 | 0.013 |
| Bond angles (°) |  |  | 1.47 | 1.70 |
| Molprobrity results |  |  |  |  |
| Ramachandran<br>outliers (%)<sup>‡</sup> |  |  | 0.1 | 0.0 |
| Ramachandran<br>favored (%)<sup>‡</sup> |  |  | 97.0 | 98.93 |
| Molprobrity<br>score<sup>‡</sup> |  |  | 1.0 | 1.30 |
| **PDB-ID** |  |  | **6FIG** | **6FIH** |

#as defined in XDS (37), <sup>†</sup>in SHARP (39), <sup>\$</sup>in Refmac5 (44), <sup>‡</sup>in Molprobrity (46)

Animal malectins and the various CBMs have been shown to bind diverse carbohydrate ligands using different surface areas of their conserved β-sandwich ‘jelly roll’ fold (27). As CrRLK1Ls have been speculated to bind cell wall components, we mapped the known carbohydrate binding sites to the ANX1 structure (28). Structural superposition of the *Xenopus laevis* malectin bound to nigerose with mal-N from ANX1 identifies a potential carbohydrate binding site on the external face of the N-terminal ANX1 β-sandwich domain (Fig. 3*A*). While the overall arrangement of secondary structure elements in this regions is conserved among *Xenopus laevis* malectin and ANX1, the protruding loops and amino-acids involved in carbohydrate binding are not present in the plant receptor ectodomain (Fig. 3*B*) (19, 29). Consistently, we failed to detect interaction of the ANX1 ectodomain with several glucose-derived disaccharides, which were previously shown to bind *Xenopus laevis* malectin domain with micromolar affinity (Fig. 3*C*). We next explored other potential carbohydrate binding surfaces: Structural alignment of the bacterial CBM22 (DALI Z-score 8.6) with the ANX1 mal-N brings the xylotetraose ligand of CBM22 in close proximity to the distal face of the β-sandwich domain normally used by B type CBMs for recognition of carbohydrate polymers. In ANX1 however, this surface area looks radically different from the known B type CBMs (Fig. 3*A*) (25, 27). Interestingly, the tetraose ligand maps to a cleft formed at the interface of mal-N and mal-C when CBM22 is superimposed onto mal-C (Fig. 3*A*). We thus tested binding of ANX1 to various cell wall derived carbohydrates of different nature and length (30, 31). However, we could not detect binding to any of the tested, commercially available cell wall polymers in isothermal titration calorimetry (ITC) assays (Fig. 3*C*). Taken together, while the ANX1/2 malectin-like domains share extensive structural homology with both animal and bacterial carbohydrate binding modules, the surface areas normally used to mediate the interaction with carbohydrate ligands appear not to be present in ANX1/2.

**Figure 3.**
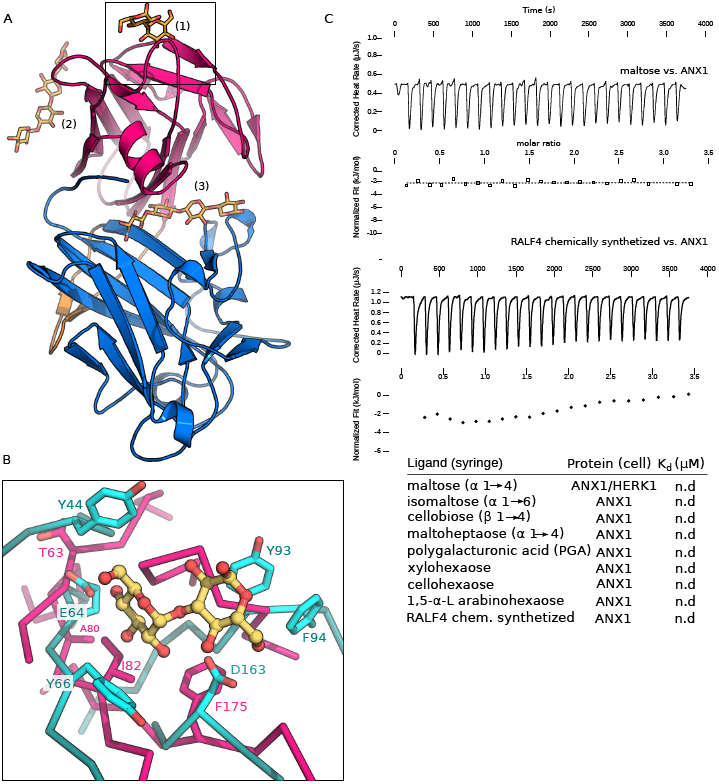
The ANX1 malectin-like domains do not form canonical carbohydrate binding sites. *A*, structural superposition of carbohydrate binding sites from the malectin protein from *Xenopus laevis* (PDB:2K46) (25) and CBM22 from *Paenibacillus barcinonensis* (PDB: 4XUR) *(26)* onto ANX1 mal-N and mal-C (colored as in Fig. 1). Carbohydrates are shown in bonds representation (in yellow): (1) The *Xenopus laevis* malectin nigerose binding site maps to the upper side of mal-N. (2) The binding surface of the xylotetraose in CBM22 is absent in mal-N, however it maps to a potential binding cleft of ANX1 when superimposed to the mal-C (3). *B*, close-up view of the *Xenopus laevis* malectin nigerose binding pocket (shown in cyan) superimposed to the ANX1 mal-N (in magenta). The residues involved in the nigerose binding are shown in cyan (in bonds representation), the corresponding residues in ANX1 (in magenta) are not conserved. *C*, isothermal titration calorimetry of D-maltose and RALF4 synthetic peptide vs. the ANX1 ectodomain and table summaries for different carbohydrate polymers (K_d_, equilibrium dissociation constant, n.d, no detectable binding).

We next analyzed the glycosylation patterns of ANX1 and ANX2. We located three and four well-defined N-glycans in our high-resolution ANX1 and ANX2 structures, respectively. In ANX1, glycans are positioned at Asn132 in mal-N and at Asn292 and 302 in mal-C (Figs. 4*A*, 5*C*). The glycosylation pattern is conserved among ANX1 and ANX2, which harbors an additional site located at Asn331. The glycosylation sites in our ANX1 and ANX2 structures map to the side and back of the tandem-malectin assembly, leaving the mal-N – mal-C domain interface and surrounding loop regions accessible for potential interactions with ligands (Fig.4*A*). In line with this, analysis of the crystallographic temperature factors in ANX1 crystals reveal that several loop regions in mal-N are rather mobile (Fig. 4*B*). It is of note that other carbohydrate binding modules also make use of large flexible loops to form specific binding sites for carbohydrate ligands (27).

**Figure 4.**
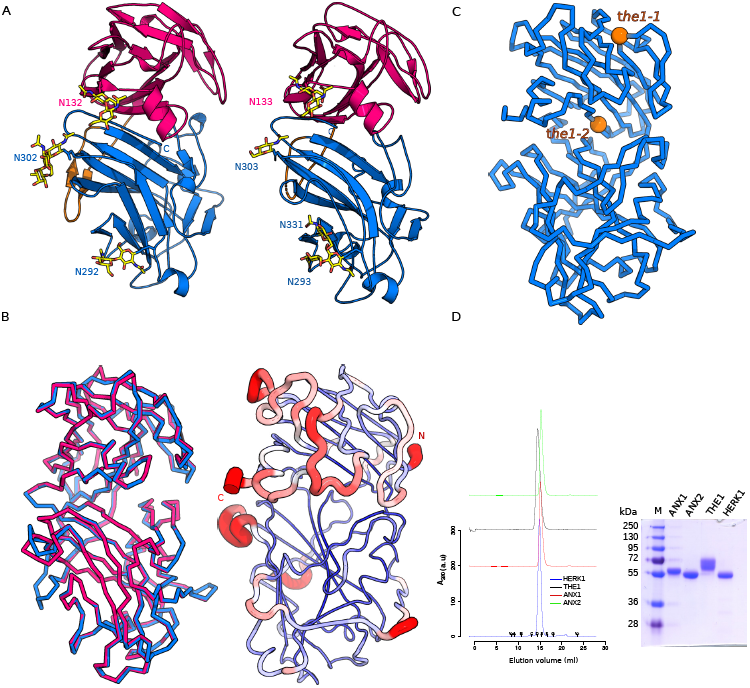
ANX1/2 share a common ectodomain architecture. *A*, ribbon diagrams of ANX1 (left) and ANX2 (right) show a strong degree of structural conservation (r.m.s.d.’s are ∼0.6 Å comparing 375 corresponding C_α_ atoms between ANX1 and ANX2) with similar orientation of their mal-N and mal-C domains (colors are as in Fig. 1). The N-glycan structures observed in ANX1 and ANX2 are highlighted in yellow(in bonds representation). *B*, structural superposition of the ANX1 (Cα trace, magenta) and ANX2 (blue) ectodomains (right) and ribbon diagram of the ANX1 ectodomain with Cα atoms colored according to their crystallographic temperature factors, from blue to red. Note that the N‐ and C-termini as well as several loop structures assembled around the ‘cleft’ region appear flexible. *C*, corresponding *the1-1* and *the1-2* alleles (shown in orange spheres) are mapped into the ANX1 structure. *D*, analytical size exclusion chromatography reveals that the ANX1 extracellular domain elutes as a monomer (red line), as do the isolated THE1 (black line), HERK1 (blue line) and ANX2 (green line) ectodomains. Void (V_0_) volume and total volume (V_t_) are shown, together with elution volumes for molecular mass standards (A, Thyroglobulin, 669,000 Da; B, Ferritin, 440,00 Da, C, Aldolase, 158,000 Da; D, Conalbumin, 75,000 Da; E, Ovalbumin, 44,000 Da; F, Carbonic anhydrase, 29,000 Da; G, Ribonuclease A, 13,700 Da.). The molecular masses of purified ANX1, THE1 and HERK1 ectodomains analysed by MS-MALDI-TOF are 58 kDa, 63kDa and 53.5 kDa, respectively. An SDS-PAGE analysis of the purified ectodomains is shown alongside.

Next, using a structure based sequence alignment of ANX1 and the related CrRLK1L THE1 involved in cell wall sensing (28), we mapped the known genetic missense alleles of THE1 onto the ANX1 structure (Figs. 4*C*, 5*C*). The *the1-1* mutation (Gly37→Asp) and the *the1-2* allele (Glu150→ Lys) are conserved among CrRLK1L family members (Fig. 5*C*). The *the1-1* mutation maps to the core of the mal-N in ANX1/2, while the *the1-2* allele is located in the mal-N – mal-C domain interface (Fig. 4*C*), suggesting that both mutations may interfere with the folding or structural integrity of the THE1 ectodomain, rationalizing their loss-of-function phenotypes (28).

**Figure 5.**
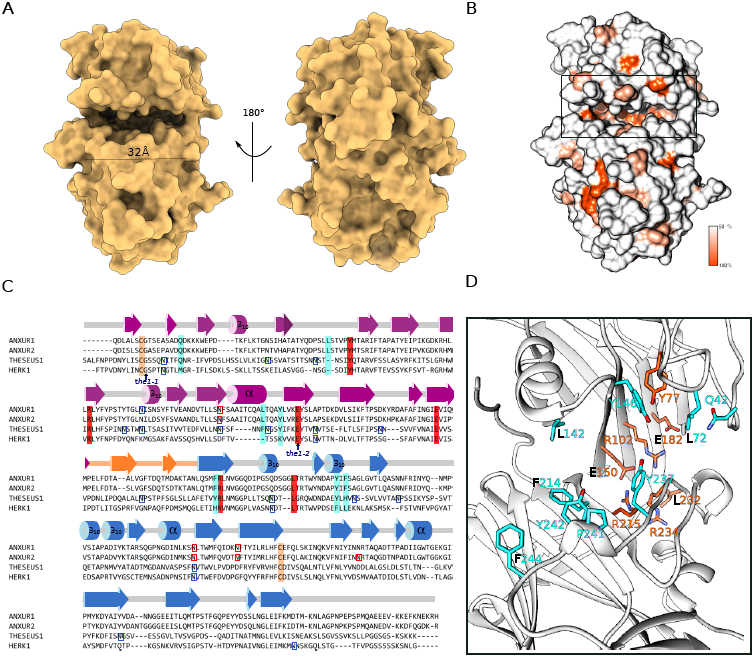
Plant tandem malectin-like receptor kinases feature a unique ligand binding cleft. *A*, front and back view of the ANX1 ectodomain in surface representation reveals the presence of a wide and deep cleft located at the interface between the N‐ and C-terminal malectin domains. *B*, surface representation of ANX1 colored according to CrRLK family sequence conservation. *C*, sequence alignment with secondary structure assignment for ANX1 calculated with the program DSSP (48) and colored according to Fig. 1. Predicted and experimentally verified N-glycosylation sites are highlighted in blue and red, respectively. The known genetic THE1 missense alleles are indicated by an arrow and the two conserved cysteine residues in CrRLK1Ls are highlighted by light orange boxes. *D*, close-up view of ANX1 (as ribbon diagram) with conserved interface residues highlighted in orange and with selected apolar and aromatic cleft-lining residues depicted in cyan (in bonds representation). Residue identifiers are according to the ANX1 sequence. Depicted residues are highlighted in *C* using the same color code.

Analysis of the lattice interactions in ANX1 and ANX2 crystals with the program PISA suggests that CrRLK1Ls ectdomains may be monomers (32) (Table 1). Consistently, we found ANX1, ANX2 and the more distantly related CrRLK1Ls HERK1 and THE1 to migrate as monomers in analytical size-exclusion chromatography experiments (Fig. 4*D*).

We next sought to identify conserved interaction surfaces in the CrRLK1L ectodomain by mapping a structure-based sequence alignment of the CrRLK1L family onto the molecular structure of ANX1 (Fig. 5). We found that many highly conserved residues (shown in orange in Fig. 5*B*) are located at the interface between mal-N and mal-C, contributing to the formation of a unique cleft structure, which is about ∼30 Å in length and provides ∼1200 Å^2^ of accessible surface area (Fig. 5*A*). Closer inspection of this cleft in ANX1 revealed the presence of several hydrophobic (Leu72, Leu142, Pro241) and aromatic amino-acids (Tyr146, Tyr214, Tyr237, Tyr242, Phe244) exposed to the solvent (Fig. 5*C, D*). Taken together our structural analysis of the ANX1 and ANX2 fertilization receptors reveal an unusual tandem arrangement of two malectin-like domains, which may have lost their ability to bind carbohydrates in previously reported CBM binding surfaces, and which form a new potential ligand binding site located at the interface of their mal-N and mal-C domains.

## DISCUSSION

Genetic studies have revealed important functions for the plant-unique CrRLK1L membrane receptor kinases in very different physiological processes, ranging from plant reproduction, cell elongation and growth to immunity (7, 23). CrRLK1Ls may regulate all these different processes by controlling specific signaling events that lead to remodeling of the cell wall. Based on their distant sequence homology with animal carbohydrate binding modules, the ectodomains of CrRLK1Ls have originally been proposed to bind carbohydrate ligands (4, 23). Our structural comparison with known animal (19, 29) and bacterial (27) carbohydrate binding modules suggests that plant CrRLK1Ls are non-canonical malectins or CBMs as they lack the conserved binding surfaces for carbohydrate ligands. These structural observations are further supported by our biochemical assays; however, our experiments cannot rule out the possibility that plant CrRLK1Ls may have evolved other, unique binding sites to sense complex plant cell wall components not commercially available for biochemical studies. Indeed, our comparative structural analysis of different CrRLK1Ls defined a cleft located at the interface between the N‐ and C-terminal malectin domains, which based on its size, surface properties and sequence conservation could represent a *bona fide* binding site for either a carbohydrate, peptide or even a protein ligand. In line with this it is of note that the plant CrRLK1L FER has been shown to interact with secreted peptide hormones of the RALF family. Direct binding of RALF1 and RALF23 to FER has been reported using pull-down and ITC assays, with dissociation constants in the low micromolar range (16, 22). In the case of ANX1/2 and BUPS1/2 RALF4 and 19 have been recently proposed as ligands (10). We thus tested direct interaction of the ANX1 ectodomain with a synthetic RALF4 peptide in ITC assays, as recently done for FER – RALF23 (16), but in our hands we could not detect binding (Fig. 3*C*). This might be due to the fact that the Cys-rich RALF peptides may have to be folded for activity (33). Alternatively, recently reported co-receptors, such as the CrRLK1Ls BUPS1/2, members of the LRR-extensin protein family or receptor-like GPI anchored proteins such as LORELEI may be required for high-affinity RALF binding (10, 34, 35).

By defining the structural architecture of CrRLK1L ectodomains, our work now sets the stage to biochemically identify and characterize ligand for this important class of plant membrane receptors and to dissect their activation mechanism.

### Experimental procedures

#### Protein expression and purification

Codon optimized synthetic genes for expression in *Spodoptera frugiperda* (Invitrogen GeneArt, Germany), coding for *Arabidopsis thaliana* ANX1 (residues 1-429), ANX2 (residues 1-431), HERK1 (residues 1-405) and THE1 (residues 1-403) ectodomains were cloned into a modified pFastBac (Geneva Biotech) vector, providing a TEV (tobacco etch virus protease) cleavable C-terminal StrepII-9xHis tag. For protein expression, *Trichoplusia ni* Tnao38 cells (36)were infected with a multiplicity of infection (MOI) of 2, and incubated for 3 days at 22°C and 110 rpm. The secreted ectodomains were purified from the supernatant by sequential Ni^2+^ (HisTrap excel; GE Healthcare; equilibrated in 25 mM KP_i_ pH 7.8, 500 mM NaCl) and StrepII (Strep-Tactin Superflow high capacity; IBA; equilibrated in 25 mM Tris pH 8.0, 250 mM NaCl, 1 mM EDTA) affinity chromatography. The proteins were further purified by size-exclusion chromatography on a Superdex 200 increase 10/300 GL column (GE Healthcare), equilibrated in 20 mM sodium citrate pH 5.0, 150 mM NaCl. For crystallization and biochemical experiments, proteins were concentrated using Amicon Ultra concentrators (Millipore, MWCO 10,000). Proteins were analyzed for purity and structural integrity by SDS-PAGE and mass spectrometry. The molecular weight of the purified, heterogeneously glycosylated proteins was determined to be ∼58 kDa (ANX1 ectodomain), ∼54kDa (ANX2), ∼53.5 kDa (HERK1) and ∼63 kDa (THE1).

#### Crystallization and data collection

Crystals of the ANX1 ectodomain developed at room-temperature in hanging drops composed of 1.0 μL of protein solution (25 mg/mL) and 1.0 μl of crystallization buffer (27% [w/v] PEG 3,350, 0.1 M Hepes pH 7.5), suspended above 0.5 mL of crystallization buffer. For structure solution, ANX1 crystals were transferred into crystallization buffer supplemented with 1 mM (NH_4_)_2_PtCl_4_ for 5 h, harvested, cryoprotected in crystallization buffer containing 15 % (v/v) ethylene glycol and snap frozen in liquid nitrogen. A 2.1 Å dataset close to the Pt L-III edge (11566.3eV, f’=-16.8, f’’=12.2) was collected at beam-line PXIII of the Swiss Light Source (SLS), Villigen, CH. As structure solution by multiple wavelength anomalous dispersion (MAD) was unsuccessful, we next collected a 1.9 Å isomorphous native dataset for SIRAS (single isomorphous replacement with anomalous scattering) analysis. Finally, a non-isomorphous high resolution native dataset at 1.48 Å was recorded.

Crystals of ANX2 (20 mg/mL) grew at room temperature in 25 % (w/v) PEG 3350, 0.1M HEPES pH 7.5 and were cryoprotected by serial transfer in crystallization buffer containing 20% (v/v) ethylene glycol as final concentration and snap frozen in liquid nitrogen. A complete dataset at 1.08 Å was collected at SLS beam line PXIII. Data processing and scaling was done in XDS (version June, 2017) (37).

### Structure determination and refinement

The structure of ANX1 was solved using the SIRAS method using a Pt derivative. Derivative and native data were scaled using the program XPREP (Bruker) and 10 Pt sites were located in SHELXD (38). Site refinement and phasing was done in SHARP (39) at 3.0 Å resolution followed by NCS averaging and density modification in PHENIX.RESOLVE (40) to 1.89 Å resolution. The density modified map was used for automatic model building in BUCCANEER (41). The resulting partial model was used to generate starting phases for the program ARP/wARP7 (42), which was used for automatic model building at 1.48 Å. The model contains two molecules in the asymmetric unit and it was completed in alternating cycles of manual model correction in COOT (43) and restrained TLS refinement in REFMAC5 (44). The final ANX1 model was used to determine the structures of ANXUR2 by molecular replacement as implemented in PHASER (45). The ANX2 solution comprises one molecule in the asymmetric unit with an associated solvent content of ∼ 40%. The structure was completed in alternating cycles of manual rebuilding in COOT and restrained TLS refinement in REFMAC. Structural validation checks were done using the program MOLPROBITY (46) and structural diagrams were prepared with PYMOL (https://pymol.org/) or CHIMERA (47).

#### Isothermal titration Calorimetry (ITC)

Experiments were performed at 25°C using a Nano ITC (TA Instruments, New Castle, USA) with a 1.0 mL standard cell and a 250 μl titration syringe. Proteins were gelfiltrated into ITC buffer (20 mM sodium citrate pH 5.0, 150 mM NaCl) and carbohydrates (maltose, isomaltose, cellobiose, heptomaltose, PGA, xylohexaose, cellohexaose, 1,5-α-L arabinohexaose) were dissolved in the same buffer. A typical experiment consisted out of injecting 10μl of the carbohydrate ligand (∼600 μM) into 50 μM ANX1 or HERK1 solution in the cell at 150 s intervals. ITC data were corrected for the heat of dilution by subtracting the mixing enthalpies for titrant solution injections into protein free ITC buffer. Sugars where obtained from Sigma Aldrich or Megazyme. RALF4 binding to ANX1 was assayed by ITC using 40 μM ANX1 solution in the cell and ∼500 μM of a synthetic RALF4 peptide (obtained by linear chemical synthesis, PSL Peptide Specialty Laboratories GmbH, Heidelberg, Germany) with protein sequence: ARGRRYIGYDALKKNNVPCSRRGRSYYDCKKRRRNNPYRRGCSAITHCYRY (design according to (16). As a control the RALF4 peptide was injected in protein free buffer. Experiments were done in duplicates and data were analyzed using the NanoAnalyze program (version 3.5) as provided by the manufacturer.

#### Size exclusion chromatography

Analytical gel filtration experiments were performed using a Superdex 200 increase 10/300 GL column (GE Healthcare) pre-equilibrated in 20 mM citric acid (pH 5.0) and 100 mM NaCl. 100 μL of the isolated ANX1 (5 mg/mL), ANX2 (4.5 mg/ml), HERK1 (5 mg/mL) and THE1 (5 mg/ml) ectodomains were loaded sequentially onto the column and elution at 0.7 mL/min was monitored by ultraviolet absorbance at 280 nm. The column was calibrated with a mixture of the high molecular weight (HMW) and low molecular weight (LMW) kits from GE Healthcare. Peak fractions were analyzed by SDSPAGE.

## Acknowledgements

We thank V. Olieric for providing beam-time and the staff at beam line PXIII of the Swiss Light Source (SLS), Villigen for technical assistance during data collection. This project has received funding from the European Research Council (ERC) under the European Union’s Horizon 2020 research and innovation programme (grant agreement no. 716358), from the Swiss National Science Foundation Ambizione Program (grant no. P200P3_161534), from SNF grant no. 31003A_173101 and from the Fondation de Famille Sandoz.

